# Comparative genomics reveals diversity and taxonomic relationships among *Clostridioides difficile* phages

**DOI:** 10.1101/2024.07.10.602917

**Authors:** Mohimenul Haque Rolin, Daniyal Karim, Nurnabi Azad Jewel, Md. Tahsin Khan, Mustafizur Rahman, Arzuba Akter, Shakhinur Islam Mondal

## Abstract

*Clostridioides difficile* is associated with life-threatening antibiotic-associated diarrhea, colitis, and toxin-mediated infections. While antibiotics are the primary treatment against *C. difficile* infections, increasing resistance necessitates alternatives. Bacteriophages and bacteriophage-derived proteins such as endolysins hold promise as potential solutions. Understanding phage biology at the genomic level is crucial for their therapeutic use. We conducted a comparative genomic analysis of 44 *C. difficile* phage genomes from public databases, examining both whole genome and proteome levels and grouping them by shared protein content. Relationships within each group were observed, and core and highly conserved genes were identified. Using genome and proteome phylogeny, average nucleotide identity, and core gene identification, we proposed an updated taxonomic classification. Nine distinct clusters were identified, without any singleton. Cluster members exhibited similar genome architecture, genome sizes, GC content, number of coding sequences, presence of core genes, and high nucleotide identity. Additionally, we propose 23 new genera, three families and the elevation of currently assigned genera to subfamilies. The lytic module proteins, endolysins, and holins were also characterized, revealing four distinct endolysin organizations with diverse domain architectures. Notably, the amidase_3 and LysM domains were highly conserved and subjected to purifying selection within the *C. difficile* phage genomes. This is the first comprehensive comparative study regarding *C. difficile* phage genomes. Our study provides valuable genomic insights that add to the current understanding of the phages. Our taxonomic analysis may improve the classification scheme of *C. difficile* phages and aid in the future classification of newly isolated *C. difficile* phage genomes.

## 1. Introduction

*Clostridioides difficile* (*C. difficile*) is a gram-positive, spore-forming anaerobic bacterium (1). It is a significant causative agent of nosocomial antibiotic-associated diarrhea, particularly in developed countries. Prevalent in the mammalian gastrointestinal tract, *C. difficile* can lead to toxin-mediated *C. difficile* infections (CDI), which manifest as a spectrum of symptoms from mild diarrhea to severe conditions such as pseudomembranous colitis, with potentially fatal outcomes. In the United States alone, *C. difficile* is responsible for over 500,000 infections annually, resulting in approximately 29,000 deaths and imposing a substantial financial burden estimated at $1-3 billion (2). CDI is linked to three well-characterized toxins – toxin A, toxin B, and binary toxin. While the former two toxins are considered primary virulence factors for CDI, *C. difficile* binary toxin has been associated with increased severity of *C. difficile* infection (3,4). *C. difficile* is commonly found in the gut microbiota of healthy individuals without causing any symptoms of the disease (5,6). However, disruptions in the gut microbiota, such as those caused by antibiotic administration or malnutrition, can predispose individuals to CDI. This imbalance in microbial composition, known as dysbiosis, can elevate concentrations of *C. difficile* toxins, facilitating colonization and the formation of biofilms in the gut. These processes contribute to the progression of severe disease states associated with CDI (6,7). Furthermore, spores of *C. difficile* are resistant to antibiotics. Following cessation of antibiotic therapy, residual spores germinate in the altered gut, leading to recurrent CDI (8,9). Faecal microbiota transplant (FMT) has been promising to prevent recurrent infections, however, several risks are involved, including the potential transfer of undesired microbes and uncertainties regarding its long-term health impact (9,10). Consequently, there is an urgent need for alternative therapeutic strategies to address recurrent *C. difficile* infections beyond FMT.

Bacteriophages or phages are viruses that infect and ultimately kill bacterial hosts (11). Phages are adapted to two lifestyles – lytic and lysogenic. Strictly lytic phages are known as virulent. In contrast, temperate phages in normal conditions are lysogenic. However, they can switch to lytic cycle in times of stress. At the end of the lytic cycle, phages lyse bacterial cell walls, leading to cell death (12). Phage therapy utilizes this property of phages, which involves the administration of phages to kill target bacteria. The major advantage of phages is their high specificity towards target bacteria while sparing beneficial bacteria (11). This intriguing feature makes phages an attractive alternative to treat CDI or recurrent CDI. Different *C. difficile* phages have been assessed for their lytic potential in multiple *in vitro* assays (13–15). Phage cocktails with 3 or 4 different phages showed more lytic efficacy compared to single phage treatments. However, rebound to lysogens was a common occurrence observed in *C. difficile* phages due to their temperate nature (9). Creation of strictly lytic phages using genetic engineering approach has been attempted but the result was similar to wild type with recombinant lytic phages reverting to lysogenic state (16). Phage-derived proteins such as endolysins offer a promising alternative to using whole phages. Phages use endolysin to hydrolyze bacterial cell wall from within, at the end of their lytic cycle (17). Several *C. difficile* phage endolysins have been cloned, characterized, and assessed for their lytic efficacy *in vitro* (9,17).

With the ever-increasing availability of fully sequenced phage genomes, *in silico* studies can elucidate on phage biology, evolution, host-phage relationships, thus, aiding in identifying therapeutically important phages and phage-derived proteins. Whole genome comparative analysis has been performed on phages infecting *Mycobacterium*, *Acinetobacter*, *Staphylococcus*, *Pseudomonas*, *Salmonella*, *Vibrio cholerae*, and Alphaproteobacteria (18–25). These studies revealed vast diversity among phage population. Phage genomes were grouped into multiple clusters based on their relatedness, with those displaying no similarity to other phage genomes designated as “singletons”. Moreover, proteins were clustered into similar groups, called “phams”, while unique proteins lacking any significant relationship with other proteins were characterized as “orphams”. Using various computational approaches such as nucleotide identity, gene sharing, single gene and network-based phylogeny, taxonomic proposals were suggested in studies involving *Acinetobacter* phages (20). The comparative genomics study on *Vibrio cholerae* phages explored therapeutic potential of the *V. cholerae* phages, showcasing a systematic approach for screening therapeutic phages in cholera and other bacterial disease (24). However, to date, no comparative studies have been conducted on *C. difficile* phages. Such studies could elucidate the relationship among the *C. difficile* phages, their genomic diversity, classification, and identify potential therapeutic options.

In this study, we conducted a comprehensive analysis of all available sequenced *C. difficile* phage genomes. Genomic comparisons were carried out utilizing different computational approaches. The genomes were grouped into multiple clusters and subclusters, and each cluster was thoroughly investigated. Furthermore, we established genus (subcluster) and family-level relationships among the phages. Finally, we examined the lysis module of CD phages, elucidating the diversity of endolysin and holin proteins.

## 2. Materials & Methods

### 2.1 Genome acquisition & annotation

Complete genomes of *C. difficile* (CD) phages were obtained from the INPHARED database (last accessed 01 March 2024) (https://github.com/RyanCook94/inphared) (26). CD-HIT was used to check for any duplication of phage genomes with 1.0 identity coverage (-c) and a word size of 10 (n = 10) (27). The genomes were then re-annotated using Pharokka (version 1.3.2), with Prodigal being used for gene calling in meta mode and PHROGs database for functional annotations (28). Phage lifestyle prediction was carried out using BacPhlip and PhaTyp (29,30).

### 2.2 Genome Clustering

Phammseqs was used to assign proteins of all genomes to phage protein families or phams with pangenome option (-p) enabled. For clustering the proteins into similar groups of proteins or phams, 35% amino acid identity and 80% coverage were used (31). Based on shared phams among the genomes, CD phages were clustered using PhamClust using a clustering threshold of 0.3 (32). A tab-separated file (TSV) file, mapping genomes to phams, and translations obtained from Phammseqs’ pangenome result were used as the input. VIRIDIC was used to calculate the intergenomic similarity of the phage genomes in default parameters (33). A 70% average nucleotide identity (ANI) cutoff value was used to assign cluster members into subclusters. Comparative genome maps were generated using gggenomes (version 1.0.0) (34). All vs all protein blasts were conducted to generate links among the genomes at an evalue 1e-05.

### 2.3 Pangenome Analysis

Initially, pangenome analysis was conducted using Pirate at 30% identity (35). For identifying core genes within cluster members, 35% identity and 50% coverage were used. However, in the case of inter-cluster core gene identification, a less stringent identity threshold of 30% was used. Additionally, CoreGenes 5.0 was utilized to identify the core genes among the phage genomes using the following options – bidirectional best hit, paralogs enabled, and an e-value of 1e-05 (36).

### 2.4 Phylogeny & Taxonomy

The VICTOR web service (https://victor.dsmz.de) was used to perform whole genome-based phylogenetic analysis (37). This method classifies and creates a phylogeny of prokaryotic viruses based on their genome. All the nucleotide sequences were compared pairwise using the Genome-BLAST Distance Phylogeny (GBDP) method, which is recommended for prokaryotic viruses. The resulting intergenomic distances were used to create a balanced minimum evolution tree with branch support via FASTME. The tree was created for each of the formulas D0, D4, and D6, respectively. Branch support was inferred from 100 pseudo-bootstrap replicates each. The trees were rooted at the midpoint and visualized with ggtree. To estimate taxon boundaries at the species, genus, and family levels, the OPTSIL program was used with the recommended clustering thresholds and an F value of 0.5 (fraction of links required for cluster fusion). ViPTree server was used to generate proteomic trees of viral sequences based on genome-wide sequence similarities (38). The analysis was performed in both query mode and reference mode. dsDNA viruses with prokaryotes as hosts were selected as references. The resulting trees were visualized and annotated in iTOL (39). Metadata for manual annotation was retrieved from Virus-Host DB (40). Additionally, single-gene phylogenetic trees were constructed utilizing the ETE3 pipeline in the genome.jp server, using amino acid sequences of the corresponding genes. Alignment and phylogenetic reconstructions were performed using the function “build” of ETE3 3.1.2 as implemented on the GenomeNet (https://www.genome.jp/tools/ete/) (41). Alignment was performed with MAFFT v6.861b with the default options (42). The resulting alignment was cleaned using the gappyout algorithm of trimAl v1.4.rev6 (43). ML tree was inferred using IQ-TREE 1.5.5 ran with ModelFinder and tree reconstruction (44). Tree branches were tested by SH-like aLRT with 1000 replicates. The resulting trees were rooted at the midpoint, annotated, and visualized in iTOL.

### 2.5 Lysis Module

For extracting the endolysins encoded by the CD phages, curated sequences of “bona fide” endolysins were used as the reference database for Diamond (45). A BLASTp identity of 30% was used for extracting endolysin sequences using Diamond (46). However, this method failed to extract all endolysins encoded by the CD phages. Thus, phammseqs assigned phams that contained endolysins were also identified using “endolysin” as the keyword. A single representative sequence was included in the subsequent analyses for genomes that encode multiple endolysins of the same domain. Endolysin domains were identified using Phmmer (47). Phylogenetic analysis was carried out by methods discussed in the previous section. Similarly, phams consisting of holin sequences were also identified, using “holin” as the keyword. Domain identification and phylogenetic analysis were carried out the same way as endolysins. To examine if endolysin and holin encoding genes went through selection pressure, analyses were carried out on selected phams of endolysins and holins. In the case of both genes, stop codons were removed and aligned with the codon-aware sequence alignment program, MACSE (48). The alignment was cleaned using trimAl with the gappyout algorithm. Datamonkey web server was utilized for selection pressure analysis using the following programs-GARD was used to take recombination event into account prior to selection pressure analysis using FEL, FUBAR, MEME, and SLAC with a p-value threshold of 0.1 (49–53). Sites under positive and negative selection, detected by each model were compared and grouped in Venn diagrams using the Venn diagram tool in the Bioinformatics & Evolutionary Genomics platform (http://bioinformatics.psb.ugent.be/webtools/Venn/).

## 3. Results

### 3.1 Diversity of *C. difficile* phage genomes

As of March 1, 2024, 44 genomes of *C. difficile* (CD) phages were available in databases. These phages were found in various sources such as water, sewage slurry, hospitals, human fecal samples, and soil. All CD phages belong to the class Caudoviricetes. While their family is still unassigned, they are currently grouped into five distinct genera: *Sherbrookevirus* (n = 11), *Leicestervirus* (n = 6), *Colneyvirus* (n = 5), *Yongloolinvirus* (n = 4), and *Lubbockvirus* (n = 2). The remaining 16 phages are yet to be classified. CD phage genomes exhibit two distinct morphotypes, Siphovirus and Myovirus, and their dsDNA genomes have both linear (n = 25) and circular (n = 19) arrangements. The phage genomes exhibit significant diversity in length, ranging from 31 kb to 135 kb, with eight genomes exceeding 100 kb, while the rest measure around 44 kb. The scatterplot in Figure 1 illustrates the length distribution of the phage genomes. The GC percentage ranges from 26% to 31%, and a highly negative correlation (Pearson R = - 0.85383) exists between the GC percentage and the genome length. Conversely, there is a highly positive correlation (Pearson R = 0.98511) between the genome length and the number of coding sequences (CDS). In contrast, a slightly negative correlation (Pearson R = -0.36264) exists between the coding density and the genome length (Figure 2). Searches with tRNAscan-se and ARAGORN revealed seven tRNAs in five of eight large phage genomes, except for LIBA6276, ES-S-0173-01, and pCD1602_4. Searches against the CARD and VFDB databases were conducted to identify resistance genes in phage genomes. Only one hit was found against the VFDB database. A hypothetical protein encoded by phage phiSemix9P1 had almost ∼80% sequence identity with a CDT type toxin (cdtB), an ADP-ribosyltransferase binding component of *C. difficile CD196*. Searches against the CARD databases yielded only a few loose hits. In addition, Minced was used to identify CRISPR spacer sequences within phage genomes. Spacer sequences were identified in the following phage genomes: CDKM15, phiCD24-1, phiCDKH01, and phiSemix9P1. Table 1 summarizes the general characteristics and features of the phages.

**Figure 1:**
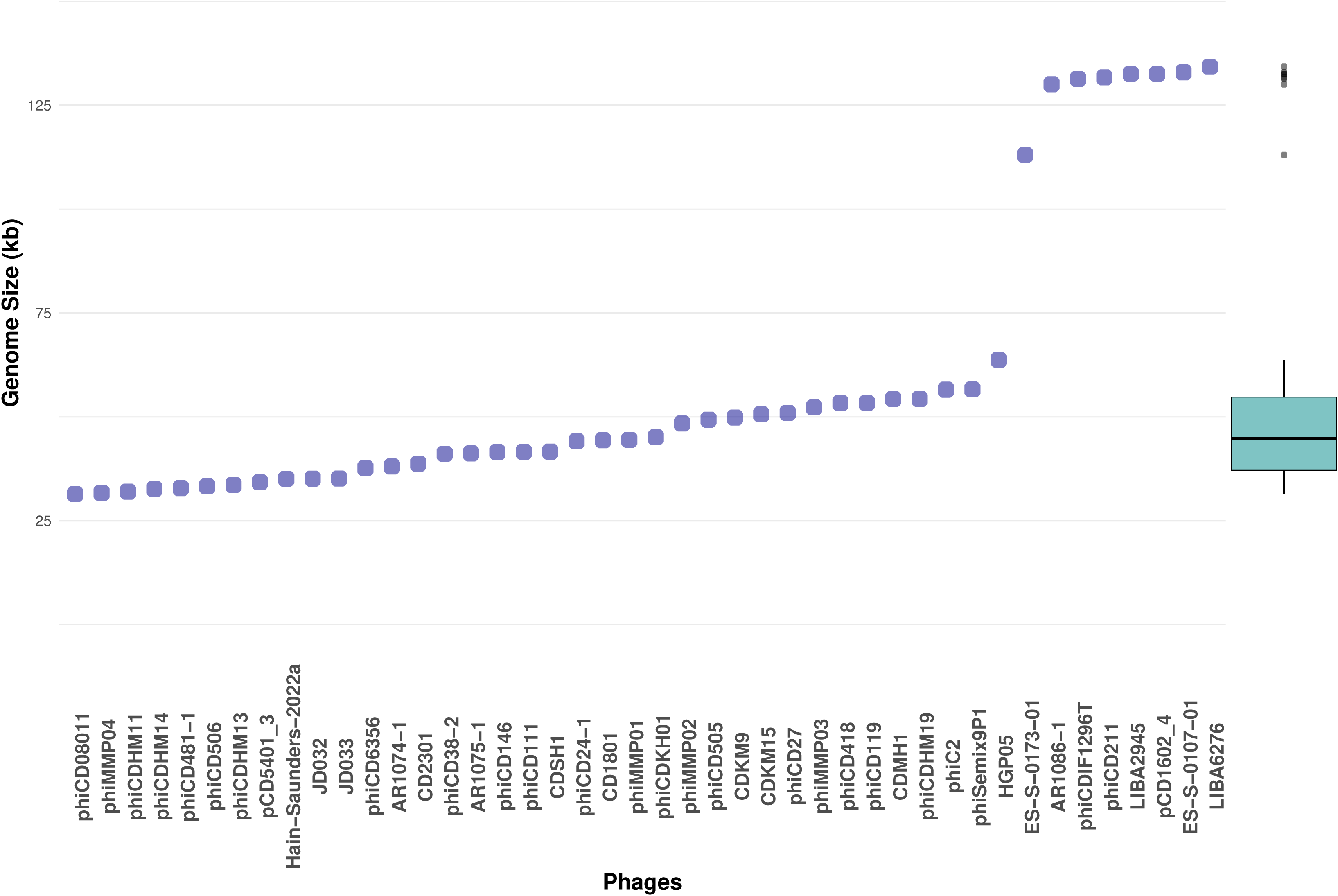
Scatterplot of genome length of the CD phages arranged in ascending order. Phages are ordered on the X-axis based on the genome length on the Y-axis. The box plot shows that most of the phage genomes are around 44 Kbp while eight genomes with >100 kbp genome length are clear outliers.

**Figure 2:**
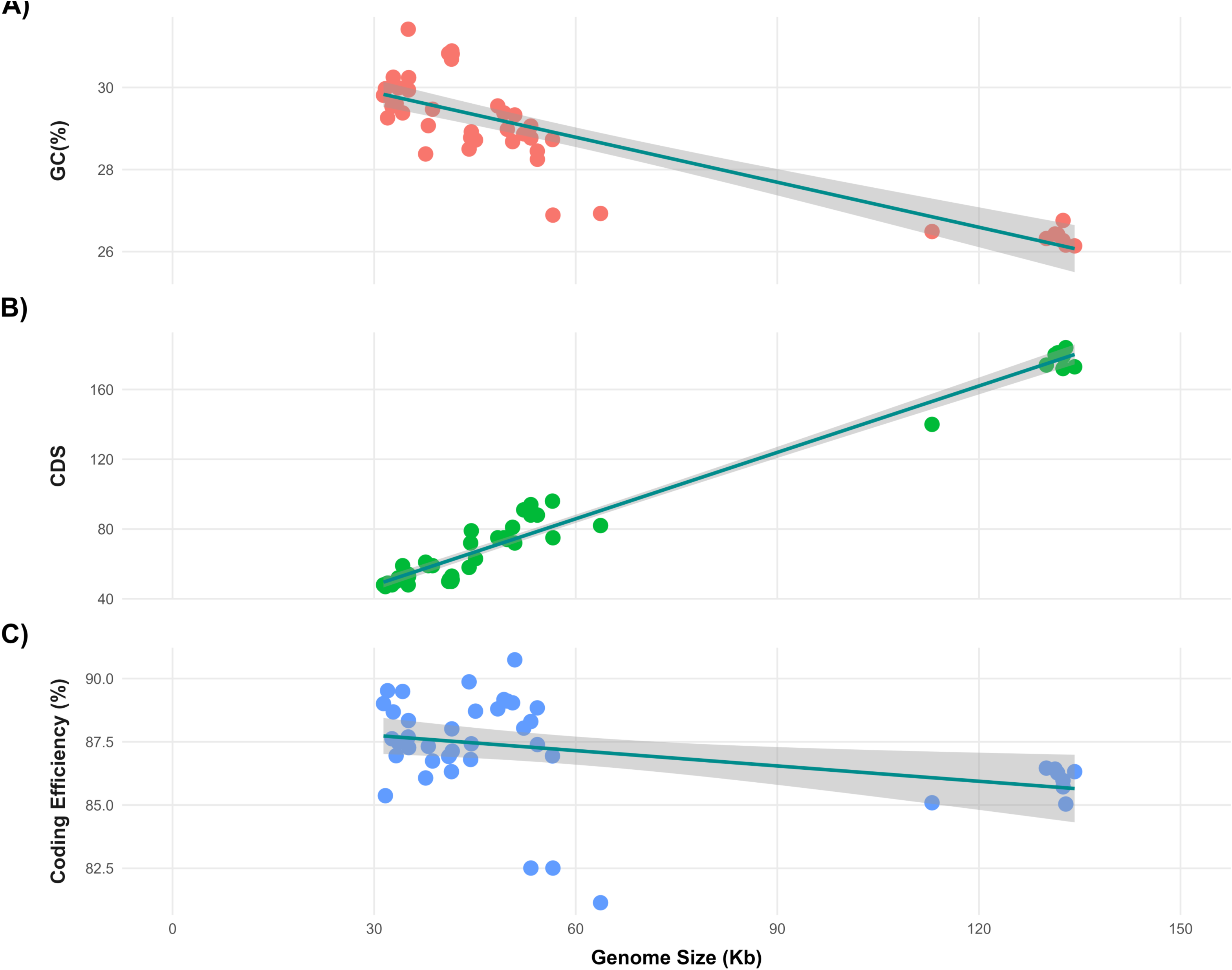
Correlation between genome size of the phages (X-axis) and three properties: GC content (%), number of coding sequences (CDS), and coding efficiency (%), each represented on the Y-axis. **A)** Correlation between genome size and GC % was highly negative (R = -0.85383) while **B)** genome size vs CDS was highly positive (R = 0.98511). **(C)** In contrast, the correlation between genome size and coding efficiency was slightly negative (R = -0.36264).

**Table 1:** General information of *C. difficile* phages.

| Accession | Phages | Length | GC | CDS | Coding_density | Genus | tRNA |
| --- | --- | --- | --- | --- | --- | --- | --- |
| MF547662.1 | LIBA6276 | 134243 | 26.14 | 173 | 86.32 | Unclassified | 0 |
| CP067347.1 | ES-S-0107-01 | 132924 | 26.16 | 184 | 85.04 | Unclassified | 2 |
| CP069348.1 | pCD1602_4 | 132519 | 26.76 | 179 | 85.72 | Unclassified | 0 |
| MF547663.1 | LIBA2945 | 132508 | 26.27 | 172 | 85.97 | Unclassified | 2 |
| NC_029048.2 | phiCD211 | 131704 | 26.42 | 181 | 86.27 | Unclassified | 1 |
| CP011970.1 | phiCDIF1296T | 131326 | 26.43 | 180 | 86.42 | Unclassified | 1 |
| OR397124.1 | AR1086-1 | 130030 | 26.32 | 174 | 86.46 | Unclassified | 1 |
| CP067352.1 | ES-S-0173-01 | 113019 | 26.49 | 140 | 85.09 | Unclassified | 0 |
| CP103976.1 | HGP05 | 63696 | 26.93 | 82 | 81.14 | Unclassified | 0 |
| KX905163.1 | phiSemix9P1 | 56606 | 26.89 | 75 | 82.51 | Unclassified | 0 |
| NC_009231.1 | phiC2 | 56538 | 28.73 | 96 | 86.94 | *Yongloolinivirus* | 0 |
| NC_028996.1 | phiCDHM19 | 54295 | 28.25 | 88 | 87.39 | *Lubbockvirus* | 0 |
| NC_024144.1 | CDMH1 | 54279 | 28.45 | 88 | 88.84 | *Yongloolinivirus* | 0 |
| NC_007917.1 | phiCD119 | 53325 | 28.77 | 94 | 82.51 | *Lubbockvirus* | 0 |
| MW512573.1 | phiCD418 | 53311 | 29.06 | 88 | 88.3 | Unclassified | 0 |
| NC_028959.1 | phiMMP03 | 52261 | 28.87 | 91 | 88.04 | *Yongloolinivirus* | 0 |
| NC_011398.1 | phiCD27 | 50930 | 29.33 | 72 | 90.74 | *Colneyvirus* | 0 |
| NC_048643.1 | CDKM15 | 50605 | 28.68 | 81 | 89.04 | *Colneyvirus* | 0 |
| NC_048642.1 | CDKM9 | 49822 | 28.98 | 74 | 89.11 | *Colneyvirus* | 0 |
| NC_028764.1 | phiCD505 | 49316 | 29.38 | 75 | 89.17 | *Colneyvirus* | 0 |
| NC_019421.1 | phiMMP02 | 48396 | 29.55 | 75 | 88.8 | *Colneyvirus* | 0 |
| MN718463.1 | phiCDKH01 | 45089 | 28.72 | 63 | 88.71 | Unclassified | 0 |
| NC_028883.1 | phiMMP01 | 44461 | 28.92 | 79 | 87.42 | *Yongloolinivirus* | 0 |
| MW512570.1 | CD1801 | 44363 | 28.78 | 72 | 86.8 | Unclassified | 0 |
| LN681534.1 | phiCD24-1 | 44129 | 28.5 | 58 | 89.87 | Unclassified | 0 |
| KU057941.1 | CDSH1 | 41619 | 30.82 | 51 | 87.13 | *Leicestervirus* | 0 |
| NC_028905.1 | phiCD111 | 41560 | 30.89 | 53 | 88.01 | *Leicestervirus* | 0 |
| NC_028958.1 | phiCD146 | 41507 | 30.69 | 50 | 86.32 | *Leicestervirus* | 0 |
| OQ703261.1 | AR1075-1 | 41195 | 30.84 | 51 | 86.95 | *Leicestervirus* | 0 |
| NC_015568.1 | phiCD38-2 | 41090 | 30.83 | 50 | 86.92 | *Leicestervirus* | 0 |
| MW512571.1 | CD2301 | 38695 | 29.47 | 59 | 86.74 | *Sherbrookevirus* | 0 |
| OR397123.1 | AR1074-1 | 38059 | 29.07 | 59 | 87.32 | Unclassified | 0 |
| NC_015262.1 | phiCD6356 | 37664 | 28.38 | 61 | 86.07 | Unclassified | 0 |
| MT193276.1 | JD033 | 35144 | 30.24 | 53 | 87.27 | *Sherbrookevirus* | 0 |
| MK473382.1 | JD032 | 35109 | 29.94 | 54 | 88.34 | *Sherbrookevirus* | 0 |
| CP103806.1 | Hain-Saunders-2022a | 35072 | 31.42 | 48 | 87.69 | *Leicestervirus* | 0 |
| CP069347.1 | pCD5401_3 | 34243 | 29.38 | 59 | 89.49 | *Sherbrookevirus* | 0 |
| NC_029116.1 | phiCDHM13 | 33596 | 29.99 | 52 | 87.47 | *Sherbrookevirus* | 0 |
| NC_028838.1 | phiCD506 | 33274 | 29.61 | 50 | 86.95 | *Sherbrookevirus* | 0 |
| NC_028951.1 | phiCD481-1 | 32846 | 30.25 | 49 | 88.68 | *Sherbrookevirus* | 0 |
| NC_048665.1 | phiCDHM14 | 32651 | 29.54 | 48 | 87.62 | *Sherbrookevirus* | 0 |
| NC_029001.1 | phiCDHM11 | 32000 | 29.26 | 49 | 89.52 | *Sherbrookevirus* | 0 |
| NC_019422.1 | phiMMP04 | 31674 | 29.97 | 47 | 85.37 | *Sherbrookevirus* | 0 |
| MW512572.1 | phiCD08011 | 31394 | 29.81 | 48 | 89.01 | *Sherbrookevirus* | 0 |

### 3.2 Genomic comparison reveals nine clusters without any singleton

For clustering phage genomes, we used protein sharing as the main criterion. PhamClust computes a Proteomic Equivalence Quotient (PEQ) for each phage pair by assessing the amino acid sequence identity of genes shared between them. PEQ values indicate the percentage of gene sharing – 0% value indicates no shared genes while 100% indicates all genes shared at 100% identity. The 44 phage genomes formed nine distinct clusters (A to I) without any singleton (Figure 3). Cluster-wise gene sharing percentages are included in the supplementary Table S1. The same clustering organization was also observed based on whole genome vs whole genome nucleotide identity visualized in the ANI heatmap (Supplementary figure S1). Additionally, using a 70% ANI cutoff value, 21 sub-clusters (denoted as A1, A2, B1, etc.) were identified (Figure S2-S8). The 44 CD phages encode a total of 3725 proteins. Proteins were assigned to 935 phams using Phammseqs at 35% mmseqs2 identity and 80% coverage. Among them, 312 (33.37%) were orphams, containing only one protein with no sequence similarity with other proteins. Phages phiCDIF1296T, phiCD38-2, phiMMP02, and phiMMP03 contain no orpham while phiCD119, ES-S-0173-01, pCD1602_4, ES-S-0107-01 contain high number of orphams (>20) (Figure S9).

**Figure 3:**
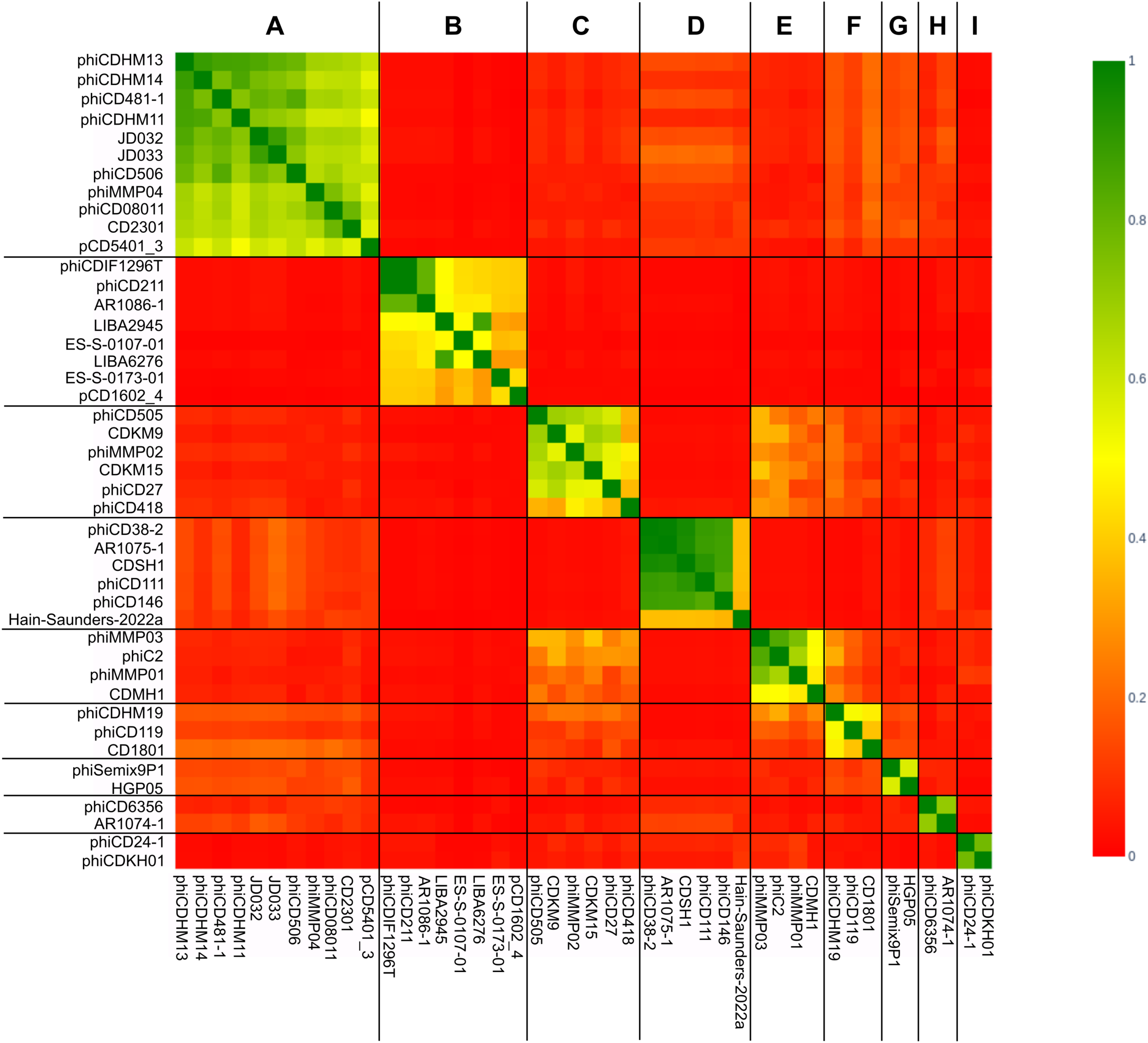
Heatmap of PEQ values among the CD phage genomes. Based on the PEQ values, 44 phage genomes were assigned to nine clusters. Assigned clusters (A to I) are shown on the top horizontal axis.

Pangenome analysis of 44 genomes using Pirate at a 30% identity threshold identified a single core gene (95% >= strains <=100%) encoding an endolysin protein. However, manual curation revealed this gene’s absence in all analyzed genomes. Further investigation identified two endolysins of the same domains present within the CD2301 phage, potentially inflating the core gene designation by Pirate. Another analysis with Phmmeseqs identified a soft-core endolysin gene (95% >= strains <=99%), further suggesting the absence of a consistent core gene. Finally, Coregenes 5.0 analysis corroborated these findings, identifying no core genes.

#### 3.2.1 Cluster A

Cluster A, the largest cluster with 11 genomes, consists of phages classified as *Sherbrookevirus* at the genus level. These phage genomes have a genome length ranging from 31 kbp to 38 kbp with a 29-30% GC percentage. Within Cluster A, the phages encode 48 to 59 genes with an average coding density of 87%. Pangenome analysis using PIRATE identified a total of 85 gene families, with 25 core genes shared among the genomes of Cluster A phages (29%). These core genes are categorized into functional groups such as head and packaging, connector, tail, DNA, RNA, nucleotide metabolism, transcription regulation, and hypothetical or unknown function. Genome comparison using gggenomes reveals a modular architecture among the phage genomes in Cluster A (Figure 4). The modular pattern followed by most of the genomes in this cluster can be described as the following - the left side of the genomes predominantly encodes structural proteins, followed by proteins involved in host lysis, while the right side contains proteins related to nucleotide metabolism, transcription regulation, integration, and excision. Cluster A genomes show high conservation across all of these modules. Notably, most phages in Cluster A encode holin except for phiMMP04, and integrase is present in all phages except for phiCDHM11. The phage genomes in Cluster A exhibit a high Average Nucleotide Identity (ANI) value ranging from 60% to 96%. Additionally, the PEQ value among Cluster A members is high as well, ranging from 50 to 90%. Cluster A genomes are divided into three distinct subclusters – A1, A2, and A3 (Figure S2). Subcluster A1 includes the sole member, pCD5401_3. Phages phiCD506, JD032, JD033, phiCD481-1, phiCDHM11, phiCDHM13, phiCDHM14 forms A2. The A3 subcluster consists of phages CD2301, phiCD08011, and phiMMP04.

**Figure 4:**
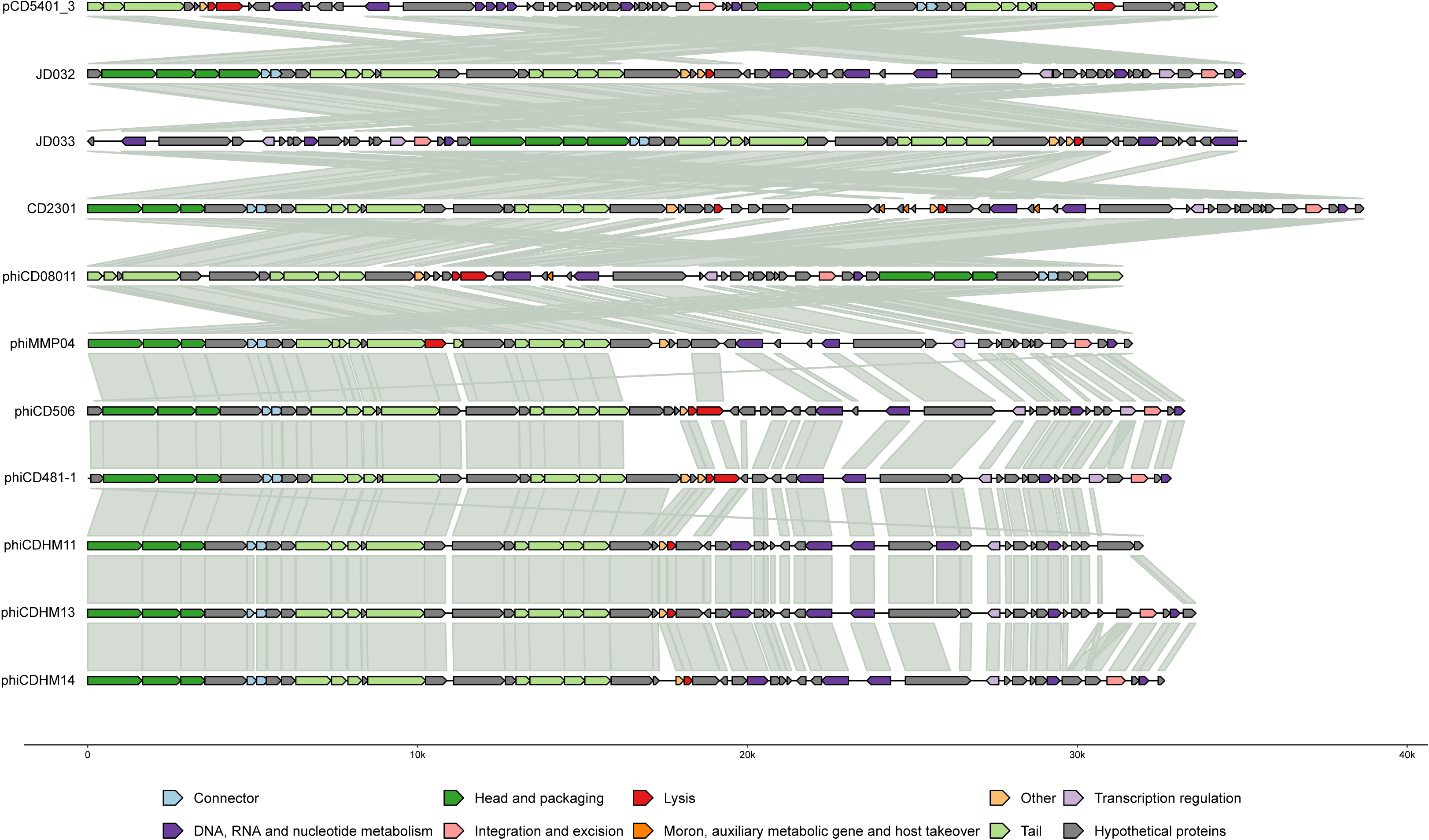
Genome comparison among cluster A phage genomes. The genomes were observed to be highly conserved. Specific modular organization is observed with high conservation visible across all modules. The genes are colored according to their functional categories as indicated in the legend.

#### 3.2.2 Cluster B

Cluster B comprises the largest eight genomes ranging from 113 kbp to 134 kbp, characterized by a lower GC content of around 26%. Following their large genome size, these phages encode a substantial number of proteins ranging from 140 to 184 with an 86% coding density. All genomes in Cluster B are yet to be classified and uniquely contain genomes with tRNA. Pangenome analysis revealed the highest number of gene families in this cluster, with 49 core genes (14%) involved in various functions like packaging and structure, DNA, RNA, nucleotide metabolism, transcription regulation, integration and excision, and lysis. A significant portion of the core genes are hypothetical proteins with unknown functions. The genome organization in Cluster B while contains specific modules like tail, lysis, and integration and excision , do not exhibit any particular pattern. Conservation is relatively low, primarily observed among structural proteins such as portal protein, virion structural protein, and a few tail proteins. Notably, integrase followed by proteins endolysin and holin that are involved in host lysis are conserved across the genomes. Synteny breaks are random and inconsistent, indicating sporadic gene transfer events. Diverse ANI values among the genomes in Cluster B are observed, ranging from 32% to 88%. Cluster B contains five subclusters (B1 to B5), three of which (B1, B2, B3) contain one member each (Figure S3). Phages LIBA2945 and LIBA6276 form B4 followed by Phages AR1086-1, phiCD211, and phiCDIF1296T forming the B5 subcluster respectively.

#### 3.2.3 Cluster C

Cluster C consists of six genomes with similar sizes ranging from 48 to 53 kbp and an average gc percentage of 29%. These genomes encode between 72 to 88 coding sequences with a relatively higher coding density. Five members of Cluster C are classified as *Colneyvirus*, while phiCD418 remains unclassified. Five subclusters are identified in cluster C - C1 to C5, with four (C1, C2, C4, and C5) containing a single member each (Figure S4). C3 is formed with phages phiCD505 and phiMMP02. Genomic comparison reveals several conserved genes among the cluster 3 genomes. These genomes collectively encode 159 gene families, with 19 core genes (12%), which is the lowest core gene count among the clusters. The core genes in this cluster are associated with functions related to the structural organization of the phage and lysis processes. These genes are confined to the left side of the genomes. Synteny breaks and random insertions-deletions (indels) are observed in the right side of the genomes, which are involved in nucleotide metabolism and host-associated functions.

#### 3.2.4 Cluster D

The highly conserved cluster D contains six genomes classified as *Leicestervirus*. Except for the phage Hain-Saunders-2022a (HS-2022a), the rest of the five phages are around 41 kbp. The genome length of Hain-Saunders-2022a is comparatively smaller, 35kbp. The genomes have a GC content of around 30-31% and encode 48 to 53 proteins with an average 87% coding density. The slight discrepancy observed in genome length in the case of Hain-Sanders-2022a is reflected in the rest of the features as well. The phages phiCD146, phiCD111, CDSH1, AR1075, and phiCD38-2 have high ANI and PEQ values (>83%), forming the subcluster D2 (Figure S5). In comparison, the ANI value of the sole member of the remaining subcluster D1, phage Hain-Saunders-2022a, is around 45-50% with the rest of the genome while PEQ values are slightly lower, around 35-37%. It is of important note that two phages AR1075-1 and phiCD38-2 are almost identical, evidenced by 100% ANI and 99% PEQ values. Cluster D genomes share 64 gene families with 33 (52%) core genes with structural, nucleotide metabolism, transcription regulation, and lysis functions.

#### 3.2.5 Clusters E and F

Cluster E represents another highly conserved group, comprising four members classified as *Yongloolinvirus*. Genome sizes within this cluster vary, with phiMMP01 having the smallest genome length at 44 kbp, while the remaining genomes range from 52 to 56 kbp. Three phages within this cluster, namely phiMMP01, phiC2, and phiMMP03, exhibit high ANI and PEQ values, ranging from 71% to 87%. Conversely, CDMH1 displays comparatively lower ANI percentages with the aforementioned phages, averaging around 53%, which is reflected in its relatively lower PEQ values. Consequently, cluster E is divided into two subclusters with phiMMP01, phiC2, and phiMMP03 forming E2 while the phage CDMH1 is the sole member of the second subcluster, E1 (Figure S6). In comparison to the other highly conserved cluster D, the core gene percentage (29%) in Cluster E is lower. Notably, among the conserved proteins, structural and hypothetical proteins are prominent.

Among the three members of Cluster F, phiCDHM19 and phiCD119 are classified as *Lubbockvirus*, while CD1801 remains unclassified. Variability in genome length is evident within this cluster, with phiCDHM19 (54 kbp) and phiCD119 (53 kbp) having larger genome sizes compared to CD1801 (44 kbp). Corresponding to the differences in genome sizes, phiCDHM19 and phiCD119 exhibit higher similarity and share high ANI and PEQ values compared to CD1801, resulting in two subclusters with CD1801 forming F1 and phiCD119, phiCDHM19 forming the subcluster, F2 (Figure S7). The genomes in Cluster F encode a total of 131 gene families, with 39 core genes (30%).

#### 3.2.6 Clusters G, H, and I

Clusters G, H, and I each consist of two unclassified members. Members of Cluster H exhibit comparatively smaller genome lengths. Notably, among all clusters, Cluster G displays the lowest coding density. Each cluster demonstrates high conservation, with ANI exceeding 60% and PEQ surpassing 57%. However, cluster G members have 61.6% ANI between them, resulting in two subclusters-G1 and G2, containing the phages HGP05 and phiSemix9P1 respectively (Figure S8). Furthermore, these clusters exhibit a high percentage of core genes: Cluster G contains 53 core genes (55%), Cluster H contains 44 core genes (62%), and Cluster 9 contains 47 core genes (72%). Of particular interest, Cluster I deviates from the typical modular pattern observed in the other clusters. Structural genes are typically located on the left side of the genome and nucleotide metabolism and integration-related genes are on the right side, a general pattern seen among the other clusters. In Cluster I, however, in the phage phiCDKH01, structural genes are present on the right side of the genome.

Genome comparison figures of clusters B to I are included in the supplementary files (Figure S10 to S14)

### 3.3 Phylogenetic analysis and taxonomic inference

Whole genome-based phylogenetic analysis using VICTOR yielded three trees with three different methods. D0, the recommended method for nucleotide analysis by VICTOR, was selected for further examination. The phylogenomic GBDP tree (Figure S15) was inferred using the formula D0 yielding an average support of 51 %. The numbers above branches are GBDP pseudo-bootstrap support values from 100 replications. The OPTSIL clustering resulted in 41 species clusters, 5 genus-level clusters, and 1 family-level cluster. However, these suggested taxonomic grouping diverged from the existing CD phage taxonomy. Therefore, the tree was also visualized and annotated in iTOL according to the current taxonomy assignment of the CD phages in the International Committee on Taxonomy of Viruses (ICTV) (Figure 5 (A)). Furthermore, the proteomic tree generated using ViPTree in query mode, including only the CD phage sequences, was similarly annotated in iTOL (Figure 5 (B)). Notably, both the genomic and proteomic trees exhibited clustering of phages from the same group. These clusters formed distinct monophyletic clades within the trees with corresponding clusters identifiable in both analyses.

**Figure 5:**
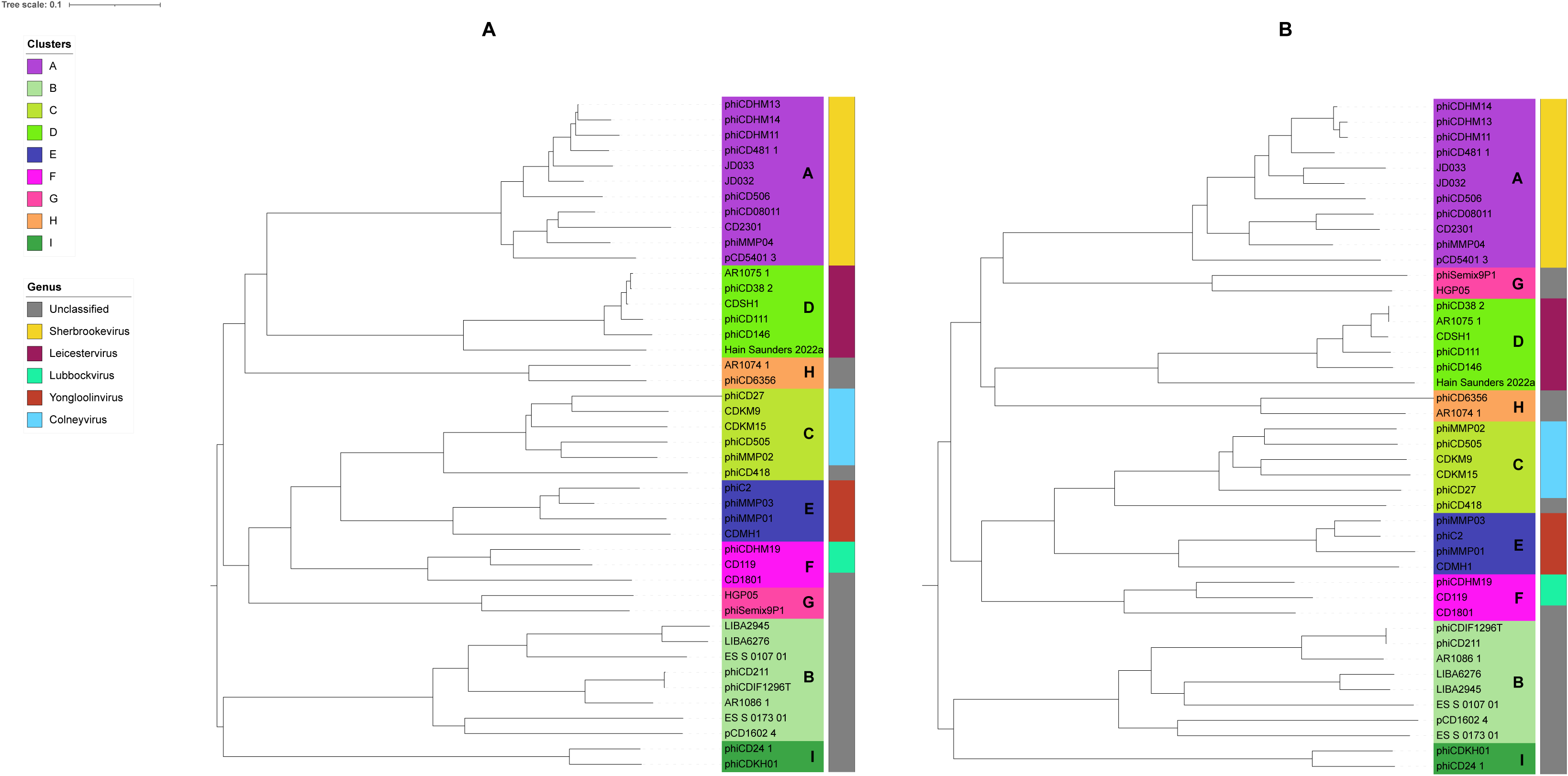
**A)** Phylogenetic tree based on whole genome generated by VICTOR. Assigned clusters and the currently assigned genus are annotated in different colors as indicated in the legend. Terminal nodes are colored according to clusters and the colored strip following the nodes represents the genus of the phages. Each cluster is distinguishable as a distinct clade within the tree. **B)** Proteomic tree generated by ViPtree using only CD phage sequences. Similarly, as the whole genome-based tree, each cluster forms a distinct clade and is discernible.

To further evaluate the cluster assignment of phages, we performed single gene phylogenetic analysis. As discussed earlier (Section 3.2), no single gene was consistently present in all the genomes. Therefore, we selected two core genes that are present in multiple clusters: portal protein and tail protein XkdN-like. Portal protein, a core gene for clusters A, B, D, E, F, H, and I, was used to construct a phylogenetic tree (Figure 6 (A)). Notably, phages from each aforementioned cluster formed distinct monophyletic clades within the tree. Interestingly, in the case of cluster C, five out of six members formed a monophyletic clade while the remaining member phiCD418 clustered with phages from cluster I. This observation highlights the mosaic nature of these phages. Similarly, the tail protein XkdN-like, a core gene for clusters C, E, F, and G, was used to generate a phylogenetic tree (Figure 6 (B)). As observed with the portal protein tree, tail proteins from phages belonging to these clusters formed separate clades. In both trees, each cluster is separately discernible, thus, solidifying cluster assignment in single-gene level.

**Figure 6:**
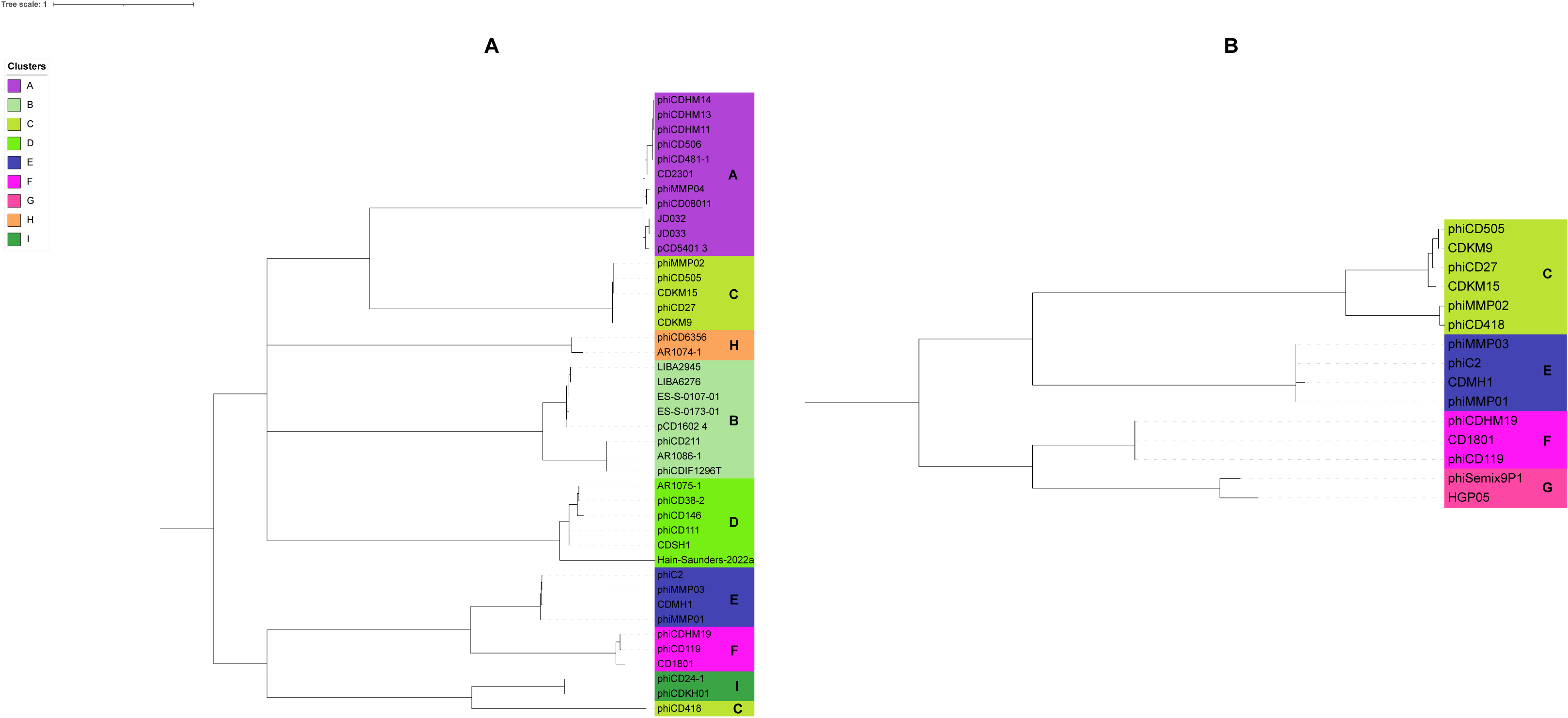
Phylogenetic tree generated with amino acid sequences of **A)** portal protein gene, **B)** tail protein XkdN-like gene. The trees were constructed utilizing the ETE3 pipeline implemented on GenomeNet (https://www.genome.jp/tools/ete/). MAFFT was used for alignment followed by alignment trimming using trimAl. The ML tree was inferred using IQ-TREE, best-fit models were LG+F+G4 (A), and LG+I (B) with branch testing by SH-like aLRT with 1000 replicates. Terminal nodes are colored according to assigned clusters indicated in the legend. Except for cluster C phage, phiCD418, portal protein genes of other phages form distinct clades with their respective cluster members. Similarly, Tail proteins of the phages form separate clades and each cluster is discernible.

The current classification system for CD phages recognizes five genera, with 28 classified phages and 16 unclassified. To establish genus-level relationships, the recommended ANI cutoff of 70% by the ICTV was employed. This analysis resulted in the subdivision of cluster members into 21 subclusters, where each subcluster represents a distinct genus. Interestingly, clusters H and I exhibited 70% ANI, indicating they each represent a single genus on their own. Consequently, this classification approach proposes the revision of CD phages into a total of 23 genera.

Furthermore, we observed that each existing genus aligns with a distinct cluster. Notably, all members within a cluster share greater than 30% nucleotide identity and possess at least 19 core genes. This high degree of sequence similarity and shared core genes suggests a sub-family level relationship among each subcluster (genus). This sub-family classification is further supported by the single gene phylogeny analysis discussed previously. Based on these findings, we propose elevating the existing genera to the subfamily level and establishing 23 novel genera for CD phages.

To investigate family-level relationships, the tree generated in reference mode considers only related (SG > 0.02) viral sequences (Figure S16). However, the lack of a significant number of shared orthologous genes, a requirement by ICTV for a single family, necessitated further analysis. We explored inter-cluster gene sharing and identified three monophyletic groups based on the recommended ICTV criteria. The first group (clusters A, D, G, and H) shared four core genes, the second group (clusters C, E, and F) shared eight core genes, and the third group (clusters B and I) shared seven core genes. Based on this analysis, we propose classifying CD phages into three families: the first comprising clusters A, D, G, and H, the second comprising clusters C, E, and F, and the third comprising clusters B and I (Table 2). The revised proteomic tree depicts the genus and family-level relationships among the corresponding clusters and cluster members (Figure 7).

**Figure 7:**
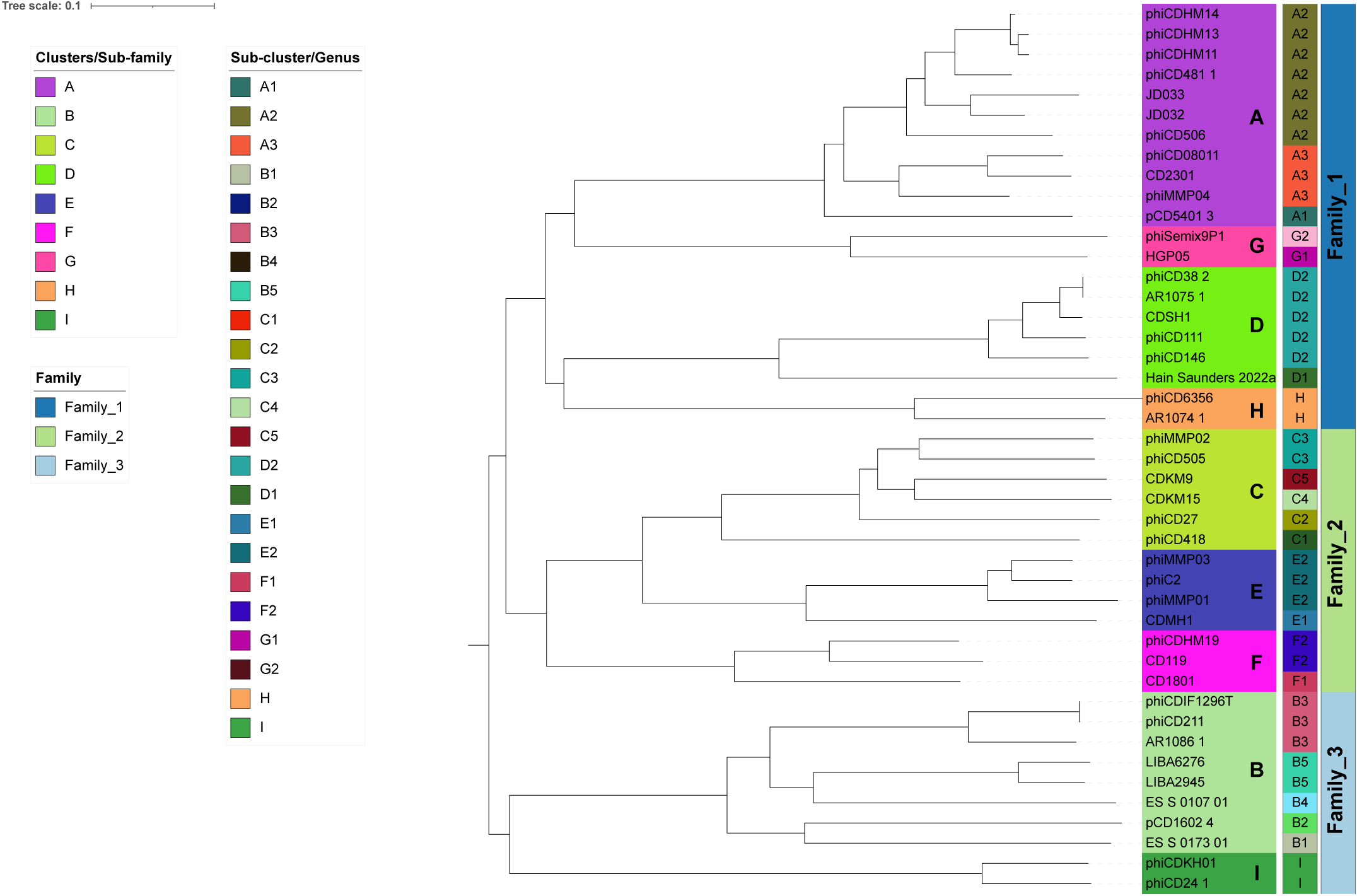
Proteomic tree annotated with updated taxonomic information. Terminal nodes are colored according to cluster/sub-family assignment. The following color strip represents sub-cluster/genus. Each sub-cluster/genus is labeled in the color strip as well. The final color strip represents family-level relationships among the clusters.

**Table 2:**
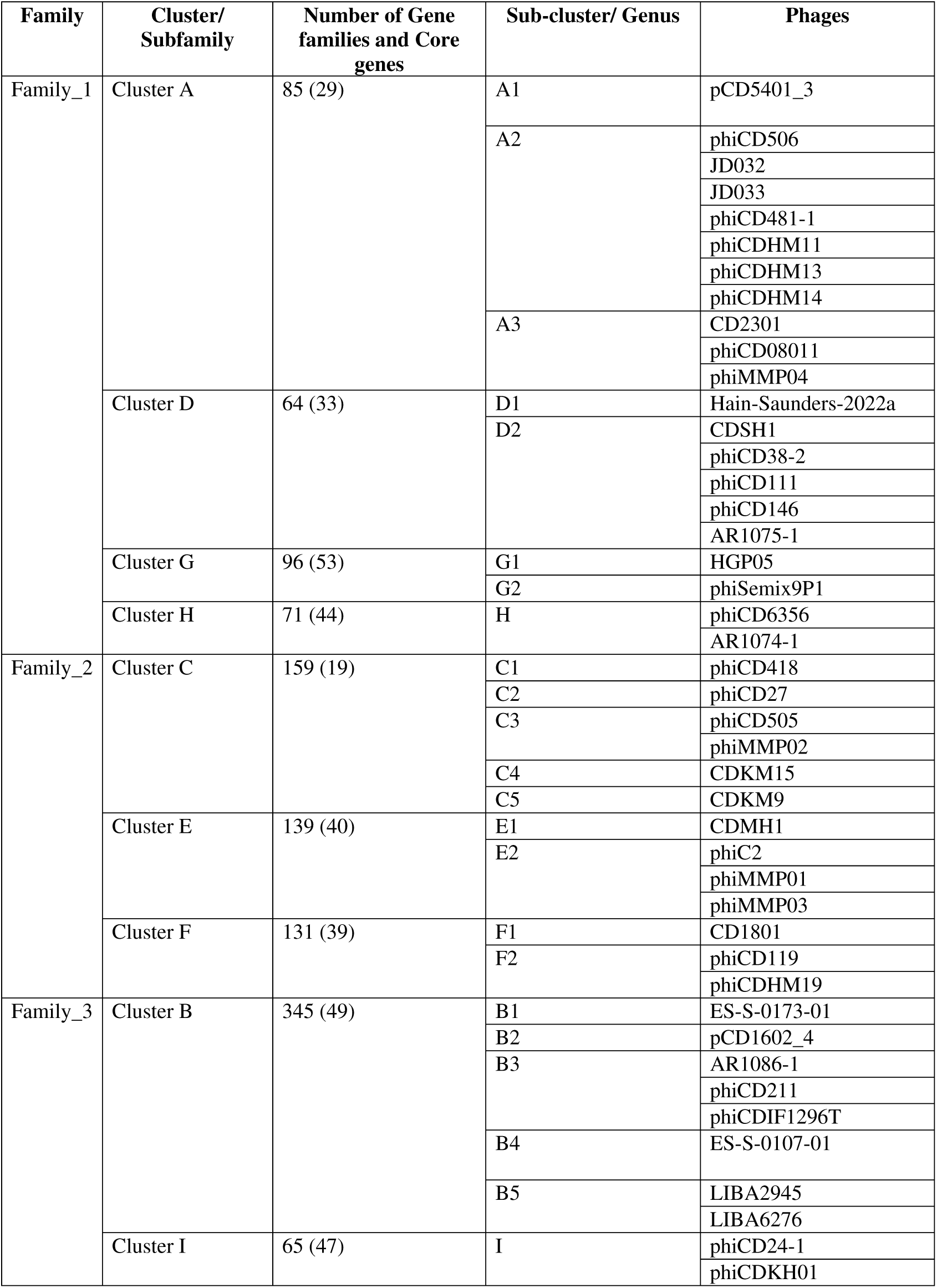
Cluster and taxonomic assignment of *C. difficile* phages.

### 3.4 Diversity of the lytic proteins

The lysis module of CD phages consists of two proteins - endolysins and holins. In this study, we identified 92 endolysins and 47 holins. Endolysins were clustered into five phams – pham_1 (n = 43), pham_12 (n = 25), pham_32 (n = 15), pham_108 (n = 8), pham_667 (n = 1). Endolysin sequences within a specific pham exhibit identical domain architecture. Pham_1, pham_32, pham_108, pham_667 contain catalytic domains amidase_3 (PF01520), NLPC-P60 (PF00877), glucosaminidase (PF01832), and amidase_2 (PF01510) respectively while endolysins in pham_12 possess the cell wall binding LysM domain (PF01476). CD phages demonstrate four distinct endolysin organizations – I) three endolysins with two encoded consecutively, while the third endolysin is encoded separately II) two endolysins encoded consecutively, III) two endolysins encoded separately, and IV) a single endolysin encoded in a genome (Figure 8). Phages in Clusters A, F, and G encode three types of endolysins containing amidase_3/amidase_2, NLPC_P60, and LysM domains, with the NLPC_P60 and LysM domain-containing endolysins being encoded by two consecutive genes. A similar arrangement is observed in Cluster B phage-derived endolysins, where two types of endolysins containing the catalytic domains glucosaminidase and amidase_3 are encoded by consecutive genes. Phages in Clusters C and E also encode two types of endolysins, with one containing the catalytic amidase_3 domain and the other containing the cell wall binding LysM domain, although they are not encoded consecutively. Conversely, phages in Clusters D, H, and I encode only one type of endolysin containing the catalytic amidase_3 domain. Endolysins of the same domain and pham form distinct clades in the phylogeny (Figure 9) except for amidase_2 containing endolysin from the orpham, pham_667, which is a singleton encoded by the CD phage phiCD481-1. Additionally, CD phage holins were clustered into three protein families-pham_5, pham_63, and pham_91. Pham_5 holins contain phage_holin_5_2 domain (PF16079) while the rest of the holin families did not yield any conserved domain when searched in phmmer. Similar to endolysins, three holin families form three distinct clades (Figure 10).

**Figure 8:**
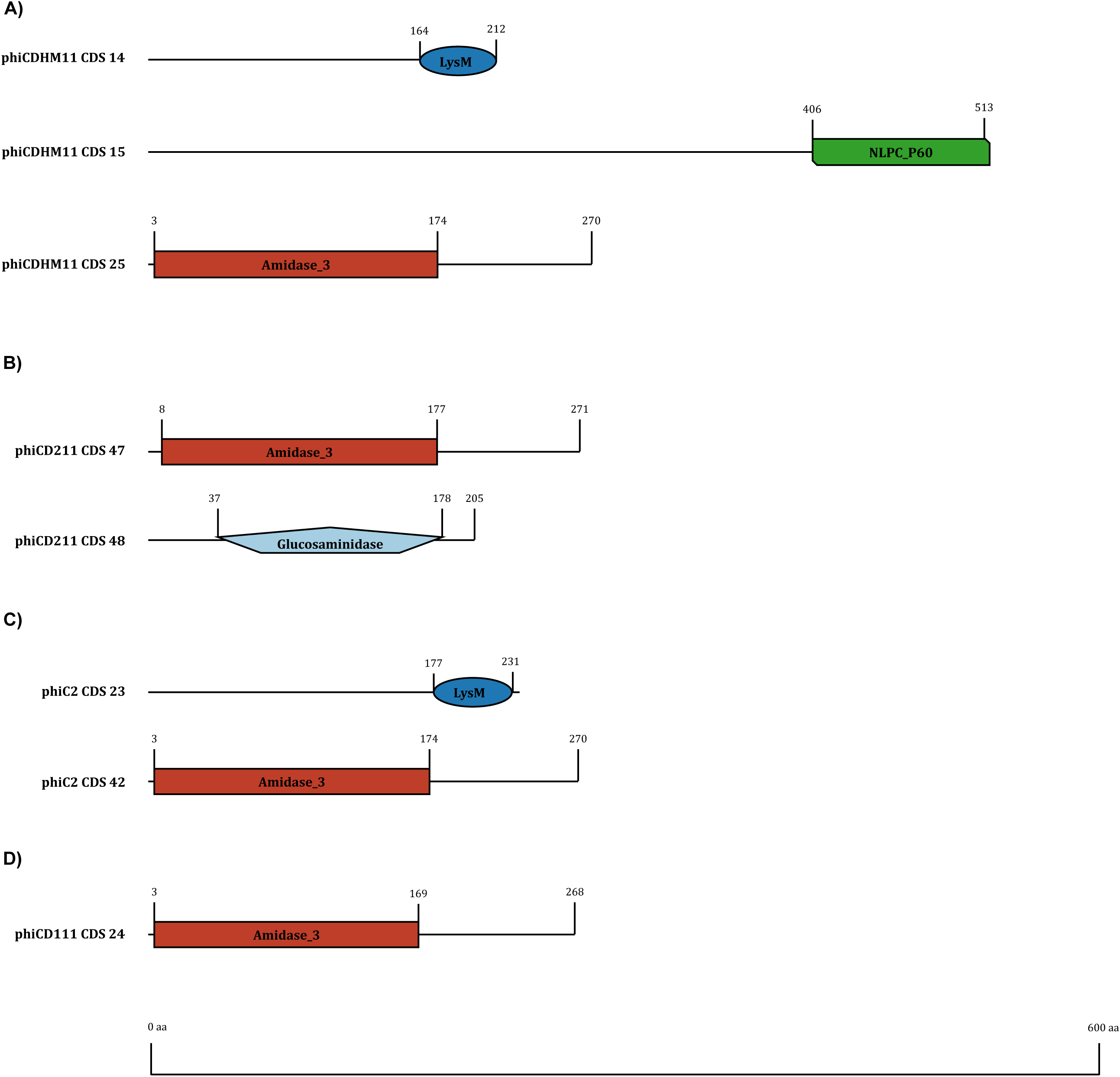
Endolysin organization and domain diversity in CD phage genomes. **A)** Three endolysins encoded in a single genome. Two are encoded consecutively, while the third endolysin is separately encoded. **B)** Two endolysins encoded consecutively encoded. **C)** Two endolysins encoded separately. **D)** Single endolysin encoded in a genome. Representative endolysins with their corresponding coding sequence numbers are included to indicate their position in the genome.

**Figure 9:**
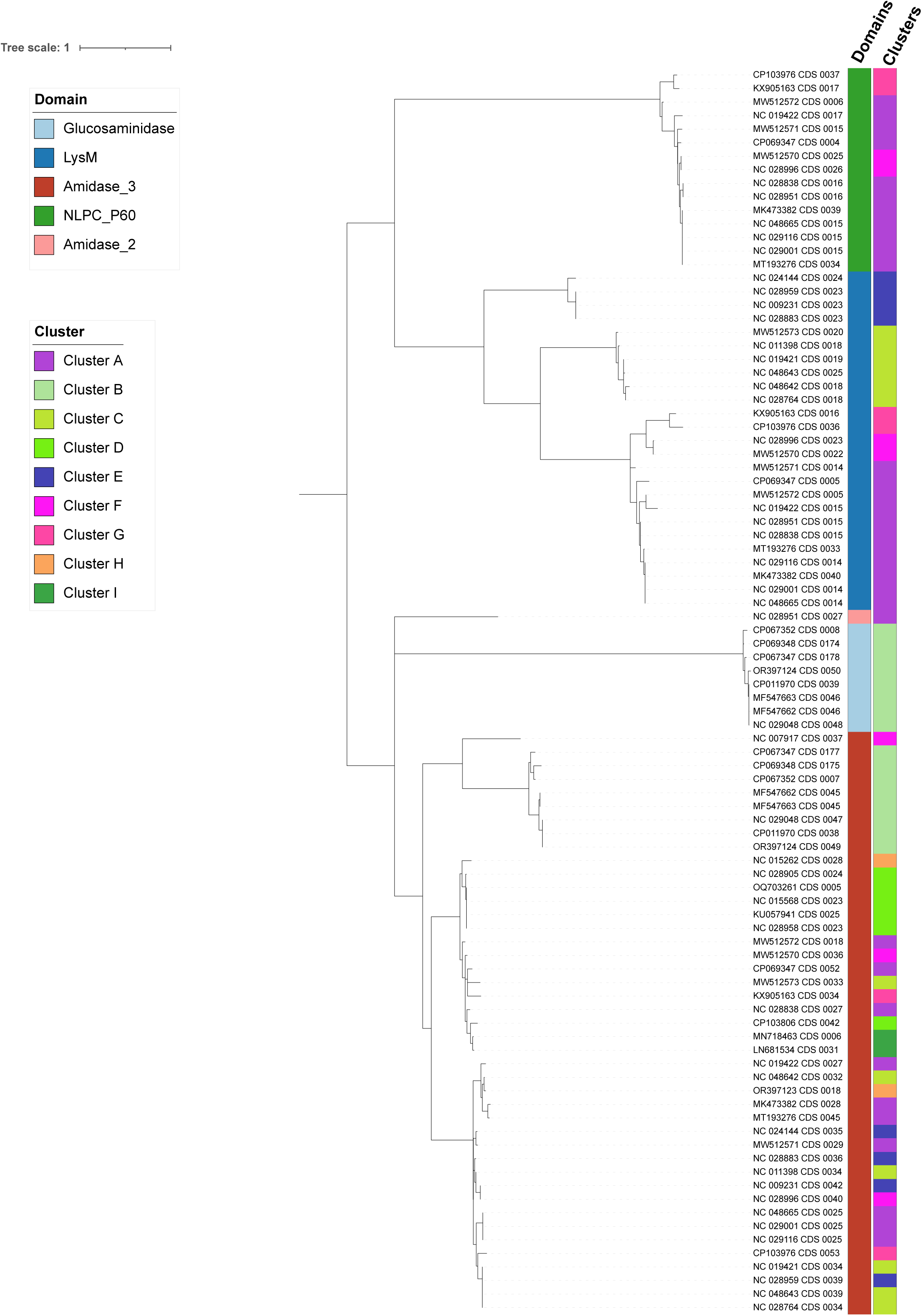
Phylogenetic tree of endolysin sequences of CD phages. The tree was constructed utilizing the ETE3 pipeline implemented on GenomeNet (https://www.genome.jp/tools/ete/). MAFFT was used for alignment followed by alignment trimming using trimAl. The ML tree was inferred using IQ-TREE, best-fit model was LG+F+G4 with branch testing by SH-like aLRT with 1000 replicates. The color strip following the terminal nodes represents domains. The second color strip represents the cluster of the corresponding host phage. Endolysins from the same domain form distinct clades, except the single amidase_2 containing endolysin. Clusters however are widely dispersed since it was observed that several single phage encodes multiple endolysins.

**Figure 10:**
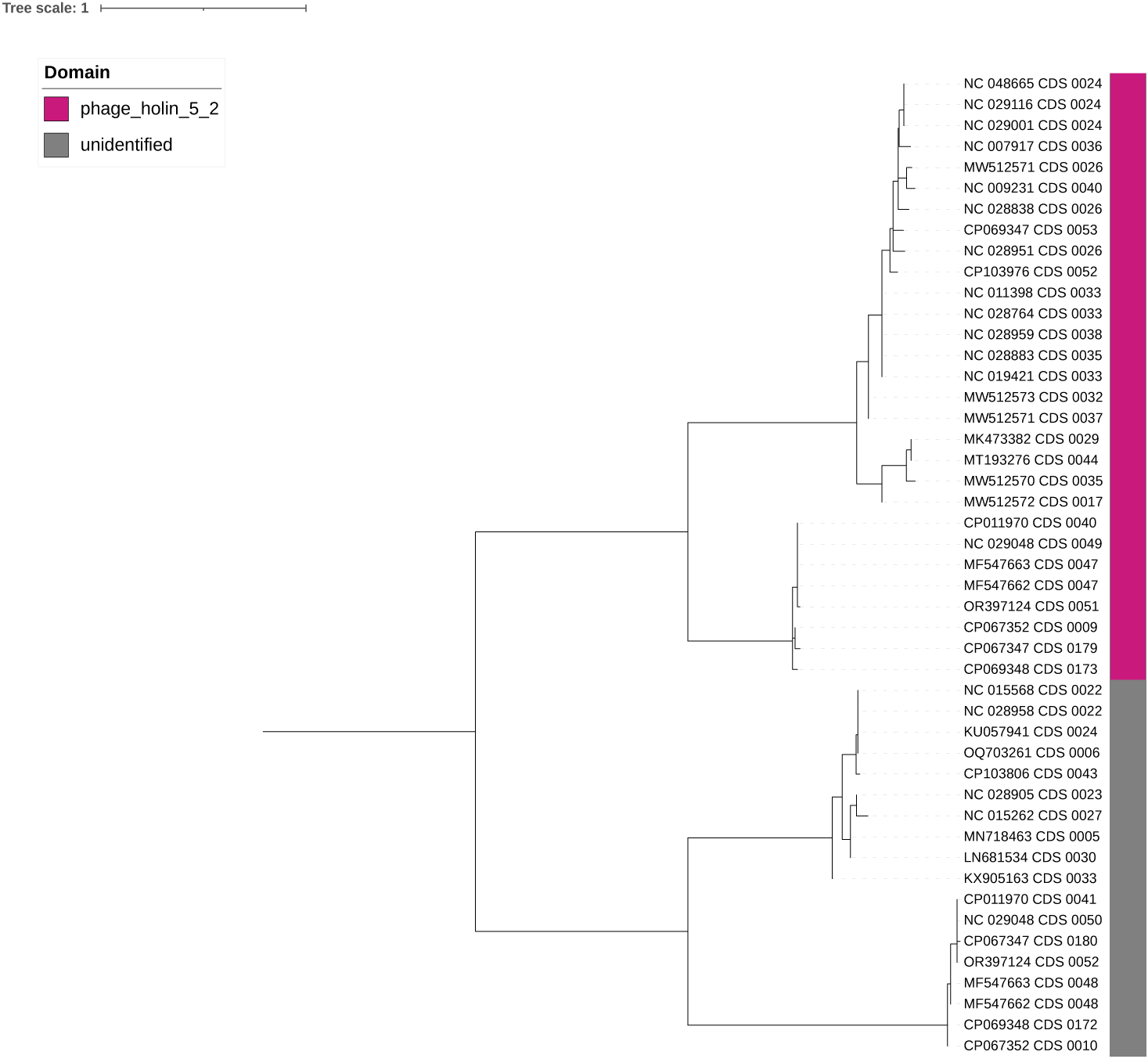
Phylogenetic tree of holin sequences of CD phages. The tree was constructed utilizing the ETE3 pipeline implemented on GenomeNet (https://www.genome.jp/tools/ete/). MAFFT was used for alignment followed by alignment trimming using trimAl. The ML tree was inferred using IQ-TREE, best-fit model was JTT+R2 with branch testing by SH-like aLRT with 1000 replicates. Holins form two major clades. One consists of phage_holin_5_2 containing holins and the other clade contains holins of unidentified domains.

Notably, most of the phages encode highly conserved amidase_3 containing endolysins (Figure S17) with 25 out of 44 phages also encoding highly conserved LysM domain-containing endolysins. This conservation is reflected in the selection pressure acting on these two endolysins. Results from Datamonkey show strong evidence of purifying selection. Only MEME result was suggestive of episodic diversifying selection at 9 of 296 sites and 7 of 211 sites for amidase_3 and LysM containing endolysins respectively. On the contrary, FUBAR suggests 182 of 296 sites, and 129 of 211 sites under pervasive purifying selection for amidase_3 and LysM respectively. In case of holins, selection pressure analysis was conducted only for pham_5 holins due to the absence of identified domains and a low number of proteins (<10) in the other phams after duplication removal. Holins, particularly those in phage_5, are subjected to strong purifying selection, as evidenced by the results of Datamonkey. Furthermore, the comparison of sites under selective pressure detected in each model is represented by Venn diagrams which illustrate the prevalence of purifying selection in CDP endolysins (Figure 11). In the case of amidase_3 containing endolysins, 92 sites under negative/purifying selection were shared by the FEL, FUBAR, and SLAC models. Five unique positive sites were detected by MEME, while three sites were shared with SLAC and FUBAR, and one site was unique to FUBAR. Similarly, for LysM-containing endolysins, 62 negative sites were detected in FEL, FUBAR, and SLAC. MEME shared one common site with FEL and FUBAR, while 27 sites were common in FEL, FUBAR, and SLAC. Supplementary Table S2 summarizes the results from Datamonkey.

**Figure 11:**
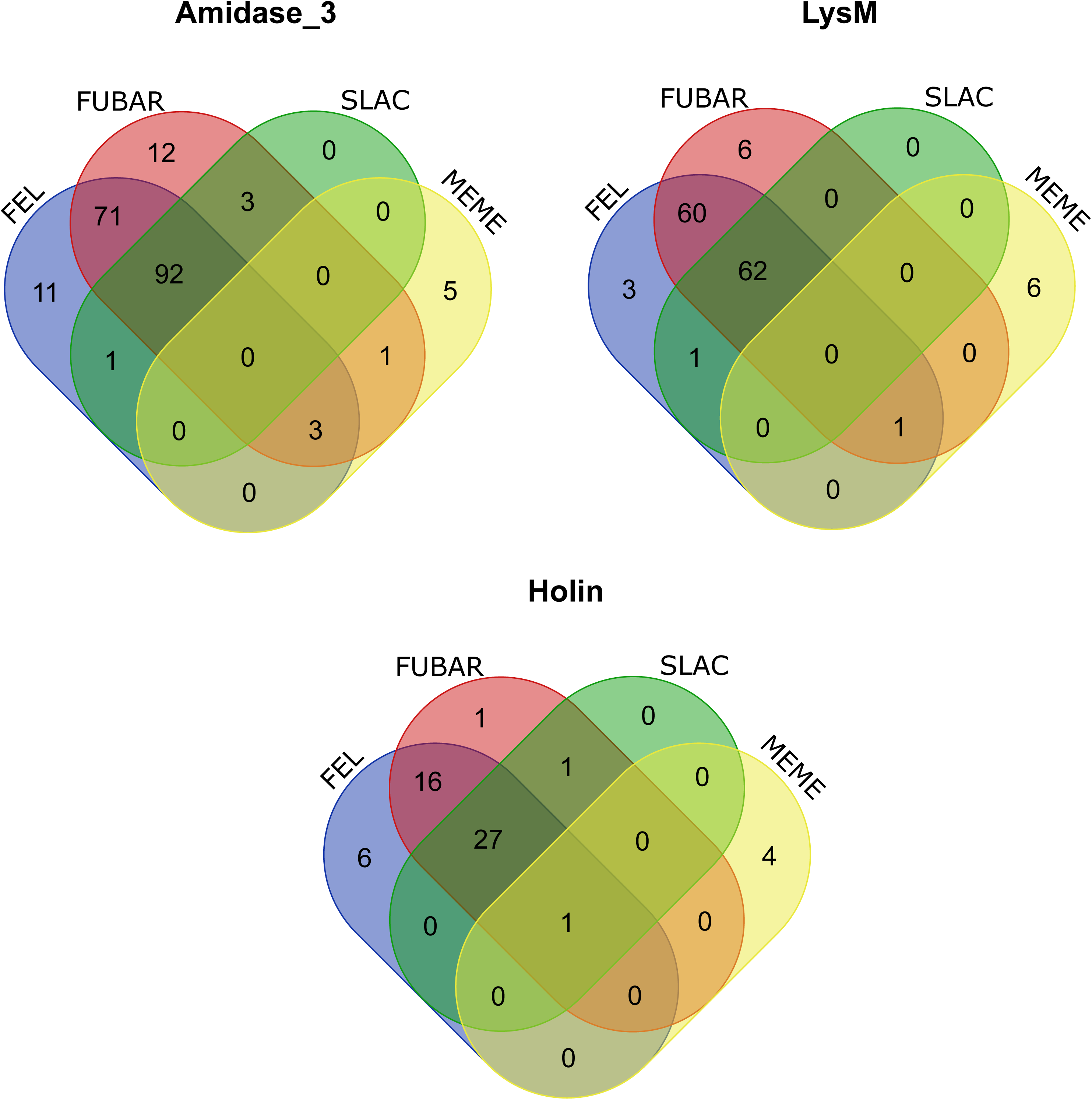
Comparison of selection sites predicted by various models in the case of amidase_3, LysM domain-containing endolysin genes, and phage_holin_5_2 domain-containing holin genes. Based on the common sites predicted by all four models FEL, FUBAR, SLAC, and MEME, negative selection is prevalent in the endolysin and holin genes.

## 4. Discussion

*Clostridioides difficile* is a major healthcare concern. However, therapeutic options, particularly for recurrent infections, remain limited. Phages and phage-derived protein endolysins offer a promising alternative due to their high target specificity. Despite this potential, our understanding of *C. difficile* phages at the genomic level is limited. This study addresses this gap by performing a comprehensive comparative genomic analysis of all available sequenced *C. difficile* phages, revealing significant insights into their diversity and taxonomy.

We observed a diverse range in genome lengths, with the number of coding sequences (CDS) corresponding proportionally to genome size. This observation aligns with established trends in phage biology. Despite this variability in genome length, CD phage genomes exhibited a low GC content within a narrow range (26 to 31%), similar to that of their host (54,55). We found that We found that larger genome sizes were associated with the presence of tRNAs, aligning with previous findings (56). This correlation was not observed in phages infecting *Pseudomonas* or *V. cholerae* where genome length did not influence tRNA presence (22,24). Computational searches for virulent genes identified a single result, the phage phiSemix9P1, which notably is the first CD phage reported to contain a binary toxin gene (57).

The clustering analysis identified nine distinct CD phage clusters, with no singletons, which is unexpected given the considerable number of singletons in other phage studies (18,22–24,58,59). This absence of singletons might change as more CD phages are sequenced. Previous studies utilized whole-genome dot plots and nucleotide identity for clustering (19,22–25,60,61); however, we emphasized protein sharing for a more accurate depiction of phage relationships, consistent with approaches used in *Gordonia* and *Staphylococcus* phages (21,62). Using a 35% amino acid identity and 80% coverage threshold as other studies (20–23,25,62), we clustered the proteins into phams. Based on the shared phams, the genomes were grouped into clusters. These clusters were supported by similar genome architecture, core gene presence, GC content, CDS number, and nucleotide similarity. Phylogenetic analyses further validated these clusters, which formed well-defined monophyletic clades. Despite less stringent criteria, no core gene was universally identified, necessitating the identification of core genes across multiple clusters for single-gene phylogeny.

The existing taxonomic classification of CD phages does not adhere to the recommended criteria for genus demarcation. By applying the ICTV-recommended 70% nucleotide identity criterion for genus demarcation (62), we propose a revision of the current genus assignments (63). Our findings suggest that the current genera should be elevated to sub-families, reassigning corresponding phages into 23 new genera. This mirrors the taxonomic structuring observed in *Acinetobacter* phage genomes, which grouped 139 phages into 8 sub-family clusters and 46 genus-level subclusters (20). It is to be noted that Clusters H and I, each with only two members and over 70% ANI between them, currently represent individual genera. However, since clusters can be interpreted as subfamilies, future discoveries of more phages in these clusters may lead to the creation of multiple genera, clarifying their subfamily-level relationships. For family-level relationships, our proteomic tree analysis indicated that CD phages form a distinct clade, separate from other phages, suggesting the need for a new family designation. This conclusion is supported by the VICTOR-assigned taxonomy (Figure S15). However, the absence of core genes or orthologues among these phages complicates the family-level assignment, a criterion emphasized by ICTV (63). We therefore explored inter-cluster relationships, identifying multiple clusters that share significant core genes, potentially representing a single family based on ICTV recommendations. The similar criteria were used to classify Lak Megaphages and Epsilon CrAss-like phages recently (64,65).

Previous studies and our computational predictions describe CD phages as lysogenic (6). This poses challenges for phage therapy due to isolation difficulties and the potential for reversion to lysogeny (16,54). Phage-encoded lytic proteins, particularly endolysins, emerge as promising alternatives for therapeutic applications (17). Our analysis of the lytic modules in CD phages revealed a diverse endolysin organization, with distinct domain arrangements. Previous studies on *Staphylococcus* phages identified four types of endolysin organization: single gene, two spliced genes, two adjacent genes, and a single gene with a secondary translational site (21). In CD phages, we observed single-gene and two adjacent-gene organizations, with separate encoding of catalytic and cell wall binding domains. Interestingly, in three CD clusters (A, F, and G), we found that two consecutive genes encode catalytic and cell wall binding domains separately, while an additional distant catalytic domain is also encoded. This suggests the possibility of these genes working together in cell wall cleavage (21). Further analysis is required to determine if the three domains function in conjunction. The catalytic amidase_3 domain and the cell wall binding LysM domain are highly conserved across CD phages, indicative of strong purifying selection, as supported by the datamonkey analysis.

*C. difficile* and its phages primarily colonize the gastrointestinal tracts of humans and animals. The dynamic environment of the gastrointestinal system likely facilitates horizontal gene transfer between hosts and phages, leading to genetic diversity. This is evidenced by the substantial proportion of orphams (33%) observed in CD phages. In *Pseudomonas* phages, orphams have significant matches in the global virome dataset (TBLASTN), indicating a largely unexplored reservoir of phages (22). Further gut and environmental metagenomics studies are expected to enhance our understanding of the diversity and gene reservoir of CD phages, potentially revealing new insights into their biology and therapeutic potential.

## 5. Conclusion

This study provides a comprehensive analysis of *C. difficile* phages, revealing their diverse characteristics, taxonomic landscape, and potential for therapeutic applications. The study identified nine distinct clusters of CD phages with varying genome sizes, GC content, and gene repertoires. Based on average nucleotide identity, protein sharing, and core gene content, the study proposes a revised taxonomic classification, elevating current genera to subfamilies and establishing 23 new genera for CD phages. Furthermore, the analysis of the lytic module suggests the presence of diverse endolysin organizations and strong purifying selection acting on catalytic and cell wall binding domains. These findings provide valuable insights into the biology of *C. difficile* phages and their potential as therapeutic agents against *C. difficile* infections.

## Acknowledgement

Not applicable.

## Ethical approval

Not required

## Supplemental materials

### Tables

**Table S1:** Cluster-wise PEQ values among the cluster members.

**Table S2:** Summary of Datamonkey results. Sites under selection inferred from various programs along with common sites detected in the programs are included in the table.

### Figures

**Figure S1:** Heatmap of ANI values among the CD phage genomes. Assigned clusters are shown on the right vertical axis. High ANI values were observed among the members of the same cluster.

**Figure S2:** Heatmap of ANI values among the members of cluster A with the subclusters highlighted.

**Figure S3:** Heatmap of ANI values among the members of cluster B with the subclusters highlighted.

**Figure S4:** Heatmap of ANI values among the members of cluster C with the subclusters highlighted.

**Figure S5:** Heatmap of ANI values among the members of cluster D with the subclusters highlighted.

**Figure S6:** Heatmap of ANI values among the members of cluster E with the subclusters highlighted.

**Figure S7:** Heatmap of ANI values among the members of cluster F with the subclusters highlighted.

**Figure S8:** Heatmap of ANI values among the members of cluster G with the subclusters highlighted.

**Figure S9:** Bar plot depicting the number of orphams per phage. Phages (X-axis) are ordered in ascending order based on the number of orphams on the Y-axis.

**Figure S10:** Genome comparison among cluster B phage genomes. The large genomes of cluster B showed the least amount of genomic conservation among the clusters.

**Figure S11:** Genome comparison among cluster C phage genomes. Despite being in the same cluster, the phage phiCD418 has relatively low similarity compared to the other five genomes, which is also observed in the figure.

**Figure S12:** Genome comparison among cluster D phage genomes. This cluster is highly conserved, especially, since the five genomes phiCD38-2, AR1075-1, CDSH1, phiCD111, and phiCD146 show very high levels of similarity which is observed in the map.

**Figure S13:** Genome comparison among cluster E **(A)** and F **(B)** phage genomes. Similar to cluster D, three genomes of cluster E are highly conserved compared to the fourth phage genome CDMH1.

**Figure S14:** Genome comparison among cluster G **(A)**, H **(B)**, and I **(C)** phage genomes.

**Figure S15:** Phylogenomic GBDP tree generated using VICTOR webserver with suggested taxa. The tree depicts 41 species-level clusters, five genus-level clusters, and one family-level cluster. Members of the same clusters are indicated by the same color. GC percentages and genome length of each phage are also represented in the tree.

**Figure S16:** Proteomic tree generated by ViPtree using CD phage and reference sequences. The inner color strip represents the viral family of the phages and the outer strip represents the phylum of their respective bacterial host. CD phages are marked as red in the tree and form a separate clade with no close relative.

**Figure S17:** Alignment of amidase_3 containing endolysins. Alignment of amidase_3 endolysins sequences was carried out with MAFFT followed by alignment trimming using trimAl. The resulting alignment was visualized with ESPript 3.0 (https://espript.ibcp.fr/ESPript/ESPript/).

